# Diet-derived vitamin B12 induces transgenerational inheritance of nematode predation through elevated vitellogenin provisioning

**DOI:** 10.1101/2025.05.21.655301

**Authors:** Shiela Pearl Quiobe, Ata Kalirad, Raphaela Zurheide, Hanh Witte, Christian Rödelsperger, Ralf J. Sommer

## Abstract

Dietary and nutritional effects can alter organismal phenotypes and influence trait inheritance over generations, so-called transgenerational epigenetic inheritance. However, the chemical nature of the stimuli and the downstream events inducing such memory remain largely elusive. The nematode *Pristionchus pacificus* exhibits mouth-form plasticity including predation and was recently shown to respond to a multigenerational *Novosphingobium* diet with the induction and subsequent transgenerational inheritance of the predatory morph. Here, we show that bacteria-derived vitamin B12 is both necessary and sufficient for transgenerational memory. Different vitamin B12 concentrations are required for the original induction and the subsequent transgenerational inheritance of the predatory morph. Mutant analysis revealed that vitamin B12 functions through methionine-synthase and involves methionine but not vitamin B9. The inherited effect acts through increased multigenerational vitellogenin transcription indicating elevated nutrient provisioning. Consistently, mutants in the vitellogenin receptor *Ppa-rme-2* are memory-defective. Thus, vitamin B12 induces vitellogenin provisioning to control organismal physiology and behavior.

---

Dietary and, more generally, nutritional effects on trait inheritance are widespread in animals including human famines. Such effects represent some of the most severe environmental influences on organismal phenotypes and can even be transmitted over multiple generations^1,2^. Indeed, a large body of evidence indicates that dietary factors can induce various morphological, physiological and behavioral changes which are often transmitted for three generations or more, so-called transgenerational epigenetic inheritance (TEI). However, the chemical nature of the inducing agents and downstream events following the original dietary stimuli are largely unknown. One major obstacle in identifying inducing stimuli from the diet and the genetic machinery underlying TEI in the receiving organism is the lack of model systems with robust phenotypes to be analyzed under laboratory conditions. However, nematodes such as *Caenorhabditis elegans* and *Pristionchus pacificus* with their short generation time, isogenic propagation and simple husbandry have started to provide mechanistic insight (for review see^3–6^).

For instance, *P. pacificus* exhibits a mouth-form dimorphism that is an example of developmental plasticity with morphological and behavioral implications and therefore, provides a robust system to analyze dietary effects and TEI.^7–9^ During postembryonic development, genetically identical *P. pacificus* worms adopt either the narrow-mouthed ‘stenostomatous’ (St) morph with a single dorsal tooth, or the wide-mouthed ‘eurystomatous’ (Eu) form with a claw-like dorsal and an opposing sub-ventral tooth (Figure 1A). Importantly, only the Eu morph allows omnivorous feeding including predation on other nematodes (Figure 1A). Mouth-form plasticity is characterized as a bi-stable developmental switch allowing the underlying gene regulatory network (GRN) to be elucidated in detail.^10–12^ Additionally, multiple environmental cues influence mouth-form development including changes in diet, establishing an easy study system for experimental manipulation.^13^

**Figure 1.**
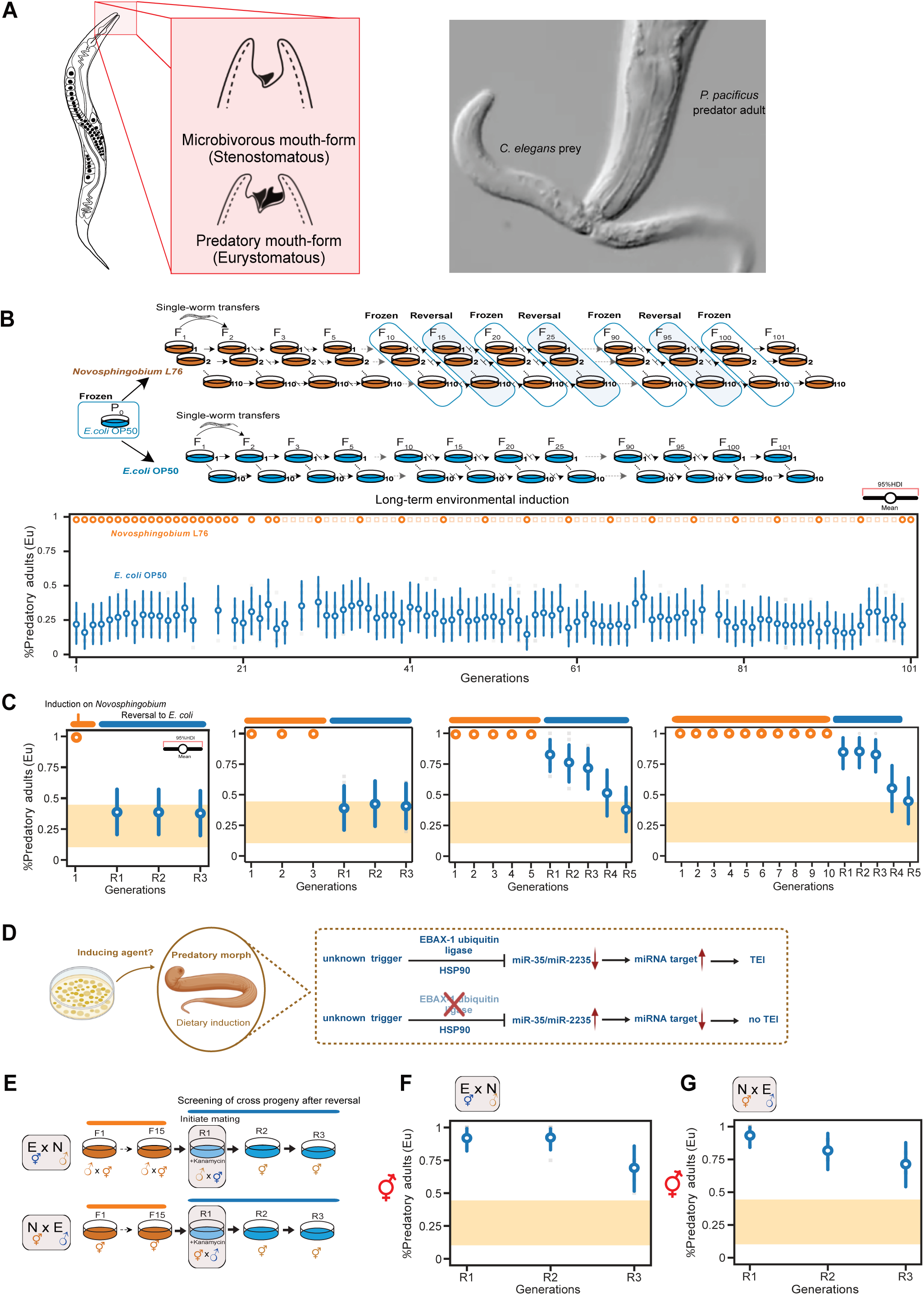
**A role for the ubiquitin ligase EBAX-1/ZWSIM8 in dietary-induced transgenerational inheritance of the predatory mouth form in *Pristionchus pacificus*** (A) Mouth dimorphism of *P. pacificus*. The eurystomatous (Eu) morph has a wide mouth with two teeth and feeds on bacteria and nematodes, whereas the stenostomatous (St) morph has a narrow mouth with a single tooth and feeds only on microbes. Right picture: Adult Eu *P. pacificus* devouring a *C. elegans* larval prey. (B) Experimental design for long-term environmental induction using distinct diets. 110 parallel lines derived from a single hermaphrodite were exposed to a *Novosphingobium* diet with 10 control lines remaining on the standard *E. coli* food source. All lines were propagated for 101 generations by single-worm descent. Periodic food reversal experiments back to the standard *E. coli* were performed all 10 generations starting in generation F15. Lower graph: Mean probability of predatory mouth-form on *Novosphingobium* (orange) and *E. coli* (blue) during the long-term environmental induction experiment for 101 generations. The y axis indicates the observed Eu mouth form frequency and the x axis, the number of generations. The 95% highest density interval (HDI) of the mean for every generation was used to visualize the probability of observing the Eu morph. (C) Transgenerational epigenetic inheritance (TEI) of the predatory mouth form after food reversal. Mean probability of predatory mouth-form after exposure to a *Novosphingobium* diet (orange) for one, three, five and 10 generations with subsequent reversal to *E. coli* (blue). A 1-generation or 3-generation exposure do not result in TEI after reversal to *E. coli*, whereas a 5-generation or 10-generation exposure results in TEI that lasts for multiple generations. (D) Model for TEI of the predatory mouth form. An unknown inducing agent results in the induction of the predatory morph, which triggers EBAX-1/ZWSIM8 activity and the degradation of clustered microRNAs of the *miR-2235/miR-35* family. (E) Experimental design for examining the contribution of male and female gametes for TEI. To determine if sperm-based transmission is necessary, *E. coli*-exposed hermaphrodites were crossed with *Novosphingobium*-exposed males. The necessity of oocyte-based transmission was observed by crossing *Novosphingobium-*exposed hermaphrodites with *E. coli*-exposed males. (F) Mean probability of predatory mouth-form in cross progenies of *E. coli*-exposed hermaphrodites and *Novosphingobium*-exposed males. (G) Mean probability of predatory mouth-form in cross progenies of *Novosphingobium-*exposed hermaphrodites and *E. coli*-exposed males (I). N*, Novosphingobium,* E, *E. coli*. See also Figure S1.

A recent study introduced a long-term environmental induction (LTEI) experiment with distinct diets by propagating 110 *P. pacificus* isogenic lines for 101 generations with associated food-reversal experiments (Figure 1B).^14^ In addition to the standard nematode laboratory food *Escherichia coli* OP50, this study used the *Pristionchus* environment-derived bacterium *Novosphingobium* L76 that was originally shown to increase killing efficiency of *P. pacificus*.^15^ The LTEI study revealed diet-derived induction of the Eu mouth form and subsequent TEI of this predatory phenotype (Figures 1B and 1C). Interestingly, TEI of the Eu morph requires a minimal exposure of five generations on the inducing diet, which is phenomenologically different from most other examples of TEI that require only a short-term stimulus (Figure 1C). Indeed, subsequent genetic studies identified a previously unknown mechanism in the regulation of TEI.^14^ Forward genetic screens for mutants defective in transgenerational inheritance indicated an essential role of the ubiquitin ligase EBAX-1/ZSWIM8 and the destabilization of clustered microRNAs of the *miR-2235/miR-35* family for transgenerational memory. Specifically, *Ppa-ebax-1* mutants show no memory whereas deletions of the *miR-35* cluster result in precocious and extended TEI of the predatory mouth form (Figure 1D). While these studies start to provide molecular mechanisms underlying TEI, nothing is known about the factors from *Novosphingobium* that induce the predatory mouth form and its subsequent TEI (Figure 1D). In this work, we studied bacterial metabolites and found that vitamin B12 is both necessary and sufficient to induce the predatory mouth form and its subsequent memory in a concentration-dependent manner. This effect is inherited through increased vitellogenin transcription and consistent with this finding, mutations in the single vitellogenin germline uptake receptor *Ppa-rme-2* are memory-defective. Thus, vitamin B12 induces vitellogenin provisioning in a multigenerational manner to control organismal physiology and behavior.

## RESULTS

### Both female and male gametes transmit TEI of the predatory mouth form

Long-term environmental induction (LTEI) is performed by switching naïve *P. pacificus* RSC011 cultures that were always grown on the standard laboratory food source *E. coli* OP50 to *Novosphingobium* L76 (Figure 1B). This *P. pacificus* strain from La Réunion Island is preferentially non-predatory (30%Eu:70%St) when grown on *E. coli*. The original LTEI experiment with 110 isogenic lines derived from the same ancestral RSC011 individual revealed dietary induction of the predatory mouth form (Figure 1B). This dietary effect is i) immediate (in generation 1 on *Novosphingobium*), ii) complete (100% Eu progeny), iii) systemic (in all 110 lines), and iv) permanent (throughout the entire experiment of 101 generations). Following the reversal from *Novosphingobium* back to the standard *E. coli* OP50 diet, we observed TEI of the predatory mouth form (Figure 1C). Importantly, such TEI is only seen after a minimal exposure to *Novosphingobium* for five generations, whereas a shorter exposure will not result in memory of the predatory morph (Figure 1C).

In theory, TEI of the Eu mouth form could be transmitted through the female gamete, the male gamete, or both. As *P. pacificus* is a self-fertilizing hermaphrodite that can outcross to males to produce cross-progeny similar to *C. elegans*, we performed crossing experiments to determine which gamete contributes to TEI of the predatory mouth form (Figure 1E). Specifically, we cultured *P. pacificus* RSC011 for 15 generations on *Novosphingobium* and performed crossing experiments with naïve *E. coli*-grown animals in all possible combinations (Figure 1E). These crosses were performed during the reversal stage on kanamycin plates and the progeny were followed for three generations to indicate TEI of the predatory mouth form. We found that the hermaphroditic progeny of a cross between *Novosphingobium*-grown males and *E. coli*-grown hermaphrodites were highly Eu for all three generations indicating TEI of the predatory mouth form (0.821 ≤HDI(θ_R1_)≤ 0.991, 0.829 ≤HDI(θ_R2_)≤ 0.99, and 0.515 ≤HDI(θ_R3_)≤ 0.859) (Figure 1F). Similarly, the progeny of *E. coli*-grown males with *Novosphingobium*-grown hermaphrodites were also highly Eu for all three generations (0.842 ≤HDI(θ_R1_)≤ 0.994, 0.672 ≤HDI(θ_R2_)≤ 0.948, and 0.542 ≤HDI(θ_R3_)≤ 0.878) (Figure 1G). In contrast, parallel crosses between hermaphrodites and males that were both *E. coli*-grown showed a low amount of Eu animals similar to naïve cultures (Figure S1). These results indicate that any gamete is sufficient to transmit the epigenetic information necessary for the TEI of the predatory mouth form.

### Vitamin B12 supplementation induces the predatory mouth form and its subsequent transgenerational memory

Next, we wanted to determine the molecular nature of the dietary stimulus inducing the predatory mouth form and its transgenerational inheritance in *P. pacificus* RSC011. The *Novosphingobium* strain used in this experimental setup was originally isolated from a *Pristionchus*-associated environment and was shown to enhance killing efficiency of the *P. pacificus* wild type strain PS312, which is naturally 100% Eu.^15^ This study also identified bacterial vitamin B12 production to cause enhanced predation, to accelerate worm development and increase brood size.^15^ Therefore, we wanted to know if vitamin B12 would also play a role in dietary induction of the predatory mouth form and TEI in *P. pacificus* RSC011 animals. For that, we transferred single naïve J4 animals to vitamin B12-supplemented agar plates with *E. coli* OP50 as food source (Fig. 2A). It is important to note that *E. coli* OP50 does not produce vitamin B12 but rather obtains its vitamin B12 from the tryptone in the agar.^15–17^ We used a final concentration of 1,000 nM of one of the two active forms of vitamin B12, methyl-Cobalamin (Me-Cbl), for supplementation, similar to previous studies. ^17^ Strikingly, we observed an immediate induction of the Eu mouth form after vitamin B12 supplementation (Figure 2B). Note that vitamin B12 supplementation results in a strong but not a complete induction of the Eu mouth form. Specifically, in three independent biological replicates out of a total of 60 assay plates we found θ^!^=0.948 in the first generation after vitamin B12 supplementation (0.878 ≤HDI(θ)≤ 0.994). This effect was seen over a period of 5 and 10 generations and was observed in all lines (Figures 2B and S2B). Thus, vitamin B12 mimics *Novosphingobium*-based induction of the predatory mouth form in *P. pacificus*.

**Figure 2.**
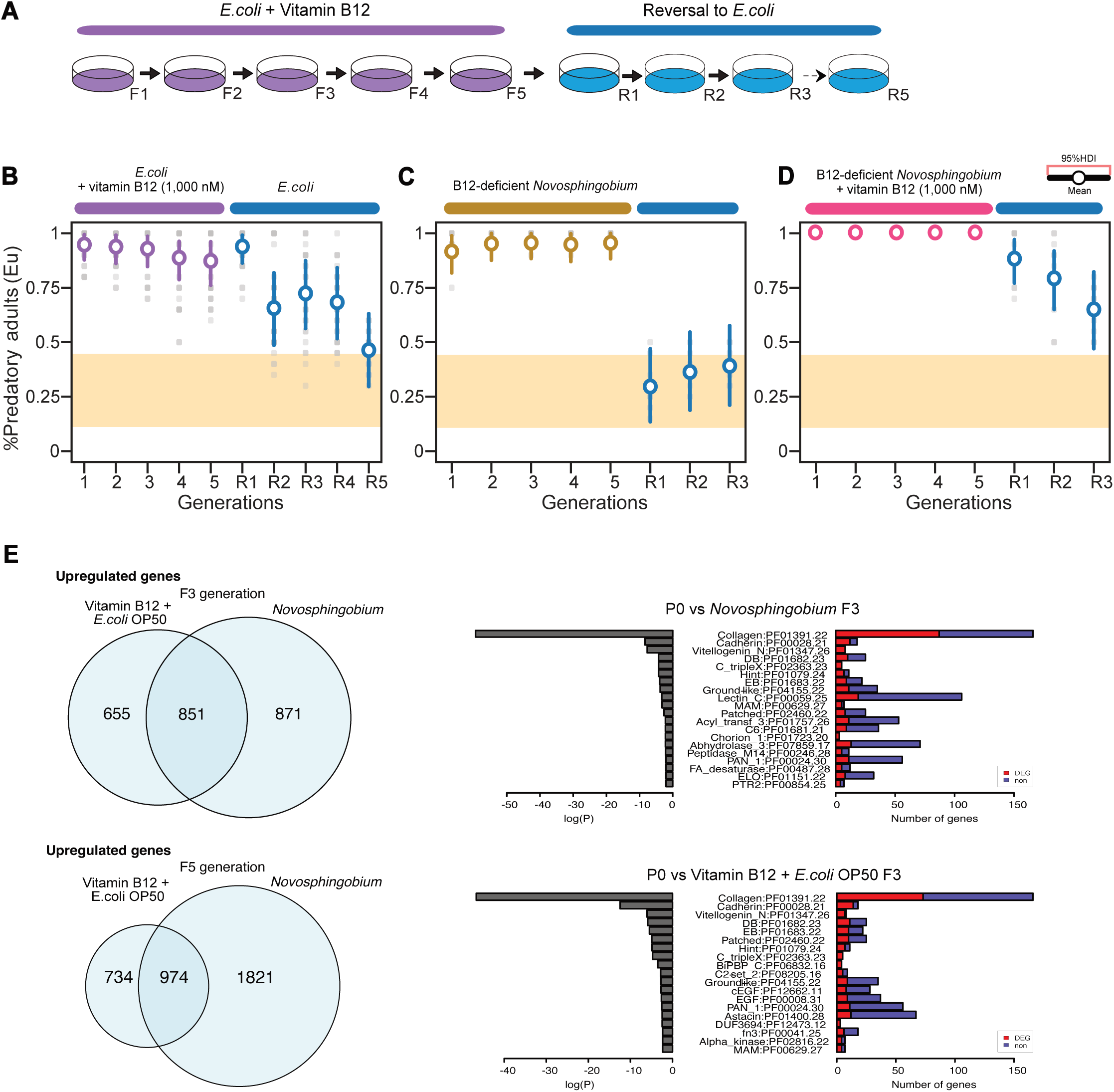
**Diet-derived vitamin B12 induces the predatory mouth form and its transgenerational inheritance** (A) Schematic diagram for vitamin B12 supplementation showing five generations on vitamin B12-supplemented plates and subsequent exposure to non-supplemented *E. coli* plates. (B) Vitamin B12 supplementation mimics induction and subsequent transgenerational inheritance of the Eu mouth form. Mean probability of predatory mouth form for five generations on 1,000 nM vitamin B12-supplemented *E. coli* and five generations on non-supplemented *E. coli*. (C) *Novosphingobium* vitamin B12-deficient mutant induces the predatory morph but not its transgenerational inheritance. Mean probability of the predatory mouth form on a *Novosphingobium* vitamin B12-deficient mutant and reversal to *E. coli*. (D) Rescue of transgenerational inheritance of the predatory mouth form by supplementing exogenous vitamin B12 to *Novosphingobium* vitamin B12-deficient mutant. Mean probability of predatory mouth form for 5 generations on vitamin B12-supplemented *Novosphingobium* mutants and subsequent reversal to *E. coli*. (E) Differential gene expression and enrichment analyses. Exposure to *Novosphingobium* and vitamin B12 supplemented *E. coli* revealed overlapping differentially-expressed transcripts relative to worm cultures grown on a standard *E. coli* diet. The pathways with most significant enrichment (FDR-corrected p value <0.01) for three generations on *Novosphingobium* and B12-supplemented *E. coli* are shown. See also Figure S2, Data S1A-D and Data S2A-D.

To determine if vitamin B12 would also induce TEI of the predatory mouth form, we established a mini-assay with a 5-generation or 10-generation exposure of worms to a vitamin B12-supplemented *E. coli* diet, followed by 5 generations on an un-supplemented diet (Figures 2A and S2A). Indeed, we observed TEI of the predatory mouth form (Figures 2B and S2B). Animals supplemented with 1,000 nM vitamin B12 showed TEI of the Eu mouth form similar to animals that were exposed to *Novosphingobium* for 5 or 10 generations. Thus, vitamin B12 supplementation mimics diet-derived induction of the predatory mouth form and its subsequent transgenerational inheritance. To our knowledge, this is the first example of vitamin-induced transgenerational memory.

### *Novosphingobium* and vitamin B12 supplementation induce overlapping transcriptomic changes

Next, we wanted to know whether vitamin B12 supplementation can elicit similar effects on gene expression as those caused by the *Novosphingobium* diet. Specifically, we examined the overlap of differential gene expression (DGE) found between vitamin B12 supplementation on *E. coli* OP50 with *Novosphingobium*-induced gene expression changes in RSC011 animals (Figure 2E). Indeed, after three and five generations of exposure, many of the *Novosphingobium*-upregulated genes were also significantly upregulated upon vitamin B12 supplementation (Figure 2E; Data S1A and S1B; Data S2A and S2B). Specifically, of the 1,506 upregulated genes after a three-generation vitamin B12 supplementation, 851 (56.5%) were also upregulated on *Novosphingobium*. Moreover, vitamin B12 supplementation for five generations resulted in 57% (974/1,708) of the upregulated transcripts being similarly induced by *Novosphingobium* after five generations (Figure 2E). The resulting upregulated transcripts from both exposures highlight major overrepresentation of protein families with the top significant signal (FDR-corrected p value <0.01) showing vast enrichment and consistent trends in Collagens, Cadherins and Vitellogenin (Figure 2E; Data S1C and S1D; Data S2C and S2D). Likewise, most *Novosphingobium*-downregulated genes largely overlap with the downregulated genes when animals were grown on *E. coli* OP50 supplemented with vitamin B12 (Figure S2D). Thus, vitamin B12 supplementation to the *E. coli* OP50 diet mimics *Novosphingobium*-mediated gene expression changes indicating that vitamin B12 is one of the major dietary metabolites influencing worm physiology under these conditions.

### Vitamin B12 is both necessary and sufficient to induce the predatory mouth form and subsequent TEI

Previous studies that established a role of dietary vitamin B12 on life history traits in *P. pacificus* and *C. elegans* were based on screening Tn5-based bacterial mutant libraries.^15,17^ We used a *Novosphingobium* vitamin B12-deficient mutant to study the effect of *Novosphingobium*-derived vitamin B12 on mouth-form plasticity and TEI of the predatory morph. Using the 5-generation mini-assay described above, we found that a *Novosphingobium* vitamin B12-deficient diet would still induce the Eu mouth form (Figure 2C). However, in contrast to a wild type *Novosphingobium* diet, the induction is not complete and plateaus ∼ 0.95% Eu (0.818 ≤HDI(θ_F1_)≤ 0.98 and 0.883 ≤HDI(θ_F5_)≤ 0.999). These results indicate that additional factor(s) besides vitamin B12 are able to induce the predatory mouth form. Thus, the complete (100%) induction of the Eu morph of *P. pacificus* RSC011 on *Novosphingobium* is due to multiple factors.

Surprisingly, we found that the *Novosphingobium* vitamin B12-deficient diet would not cause TEI of the predatory mouth form. When we reverted worm lines that were exposed to the *Novosphingobium* vitamin B12-deficient diet for 5 generations back to *E. coli*, we observed an immediate return to the base line response level (0.135 ≤HDI(θ_R1_)≤ 0.47) (Figure 2C). Similarly, when we reverted RSC011 lines that had been exposed to this diet for 10 generations, we also did not observe TEI of the predatory mouth form (Figure S2C). In contrast, vitamin B12 supplementation for 5 or 10 generations to the *Novosphingobium* vitamin B12-deficient diet restored the normal TEI response after reversal to *E. coli* OP50 (0.772 ≤HDI(θ_R1_)≤ 0.971, 0.65 ≤HDI(θ_R2_)≤ 0.919, and 0.47 ≤HDI(θ_R3_)≤ 0.832) (Figures 2D and S2D). These results indicate that vitamin B12 is both necessary and sufficient to induce the TEI of the predatory mouth form.

### Vitamin B12-induced TEI of the predatory mouth form requires *metr-1*

In animals including humans, vitamin B12 acts as a cofactor of two enzymes, methionine-synthase in the cytoplasm and methylmalonyl coenzyme A (CoA) mutase in mitochondria (Figure 3A).^18^ In *P. pacificus* as in *C. elegans*, the methionine-synthase and methylmalonyl coenzyme A mutase are encoded by the *Ppa-metr-1* and *Ppa-mce-1* genes, respectively. We targeted both genes in *P. pacificus* RSC011 by CRISPR to study a potential role in dietary induction and TEI of the predatory mouth form (Figure 3B). We obtained two frameshift alleles of *Ppa-metr-1* and *Ppa-mce-1* each, and tested all of them in our standard assay (Figure 3B). First, we studied the response of *Ppa-metr-1* mutants to an *E. coli* diet supplemented with 500 nM, 1,000 nM and 1,500 nM vitamin B12. *Ppa-metr-1* mutant animals failed to respond to vitamin B12 indicating that the role of vitamin B12 in inducing the Eu mouth form requires *Ppa-metr-1* (Figure 3C). Second, we grew *Ppa-metr-1* mutants on *Novosphingobium* and observed a mean induction of θ^!^=0.96 throughout the exposure of 5-10 generations (Figures 3D, S3A and S3B). This result is likely due to the other *Novosphingobium* factor(s) that can induce the Eu mouth form as indicated in the experiments using the *Novosphingobium* vitamin B12-deficient diet (Figures 2C and S2C). Importantly, when we reverted *Ppa-metr-1* mutants from *Novosphingobium* back to *E. coli*, we observed no TEI of the predatory mouth form (Figures 3D, S3A and S2B). These findings indicate that vitamin B12 acts as a cofactor of methionine-synthase in the transgenerational inheritance of the Eu mouth form. Third, we tested *Ppa-mce-1* mutants and observed an induction of the Eu mouth form on *Novosphingobium* similar to wild type animals (Figures 3E, S3D and S3E). After reversal, *Ppa-mce-1* mutant animals responded similar to wild type controls indicating the *Ppa*-MCE-1 is not involved in the TEI of the predatory mouth form (Figures 3E, S3D and S3E). Note however, that the response of *Ppa-mce-1* mutants to vitamin B12-supplemented *E. coli* diet only resulted in a partial Eu induction (0.256≤HDI(θ)≤ 0.616 for the highest vitamin B12 concentration) (Figure S3C). Together, our mutant analysis identifies a requirement of *Ppa-metr-1* for the vitamin B12-mediated induction of the predatory mouth form and its subsequent TEI.

**Figure 3.**
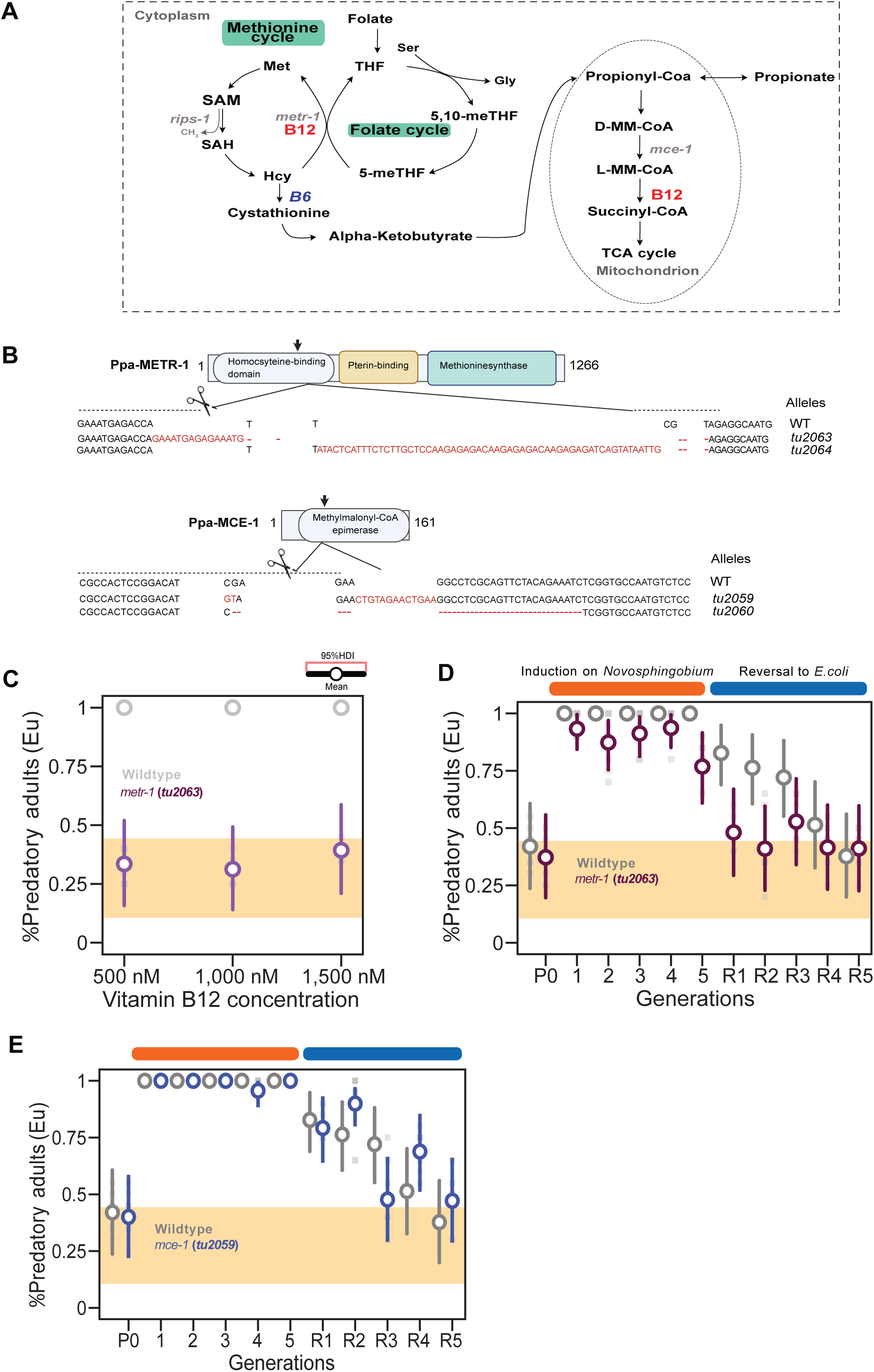
**Vitamin B12 functions as a cofactor for methionine synthase *metr-1* in the transgenerational inheritance of the Eu morph** (A) Requirement for vitamin B12 as cofactor for two enzymes in the cytosol and mitochondria. In the cytoplasm, it acts as cofactor of methionine synthase (*metr-1)* as part of the one carbon cycle. In the mitochondria, vitamin B12 is a co-factor of methylmalonyl coenzyme A mutase. Both genes (*metr-1* and *mce-1*) encoding the metabolic enzymes in which vitamin B12 acts as a cofactor are shown. Red, vitamin B12; Blue, vitamin B6. (B) CRISPR/Cas9-induced mutations in *Ppa*-METR-1 and *Ppa*-MCE-1 with target locations indicated in respective protein domains (sgRNA, arrow). Molecular lesions of isolated mutations via CRISPR/Cas9 are also shown. (C-D) Requirement of *Ppa-metr-1* in transgenerational inheritance of vitamin B12-induced Eu mouth form. (C) Mean probability of predatory mouth-form in *Ppa-metr-1* mutant animals on vitamin B12-supplemented *E. coli* compared to wild type. (D) Mean probability of the predatory mouth form in *Ppa-metr-1* mutant animals on a *Novosphingobium* diet and reversal to *E. coli* compared to the wild type response. (E) Mean probability of the predatory mouth form in *Ppa*-*mce-1* mutant animals exposed to *Novosphingobium* and reversal to *E. coli* compared to wild type response. See also Figure S3 and Table S1.

### Induction of the Eu morph and its subsequent TEI require different concentrations of vitamin B12

Given that vitamin B12 mimics the effect of *Novosphingobium* on mouth-form plasticity and the transgenerational memory of the predatory morph, we next performed proper quantifications of these effects. While our original experiments used a final concentration of 1,000 nM Me-Cbl, which corresponds to the amount used in previous studies^15,17^, we determined the minimum concentration necessary to induce these effects. When we reduced the concentration of vitamin B12 to 500 nM, we observed a strong induction of the predatory mouth form (Figure 4A).

**Figure 4.**
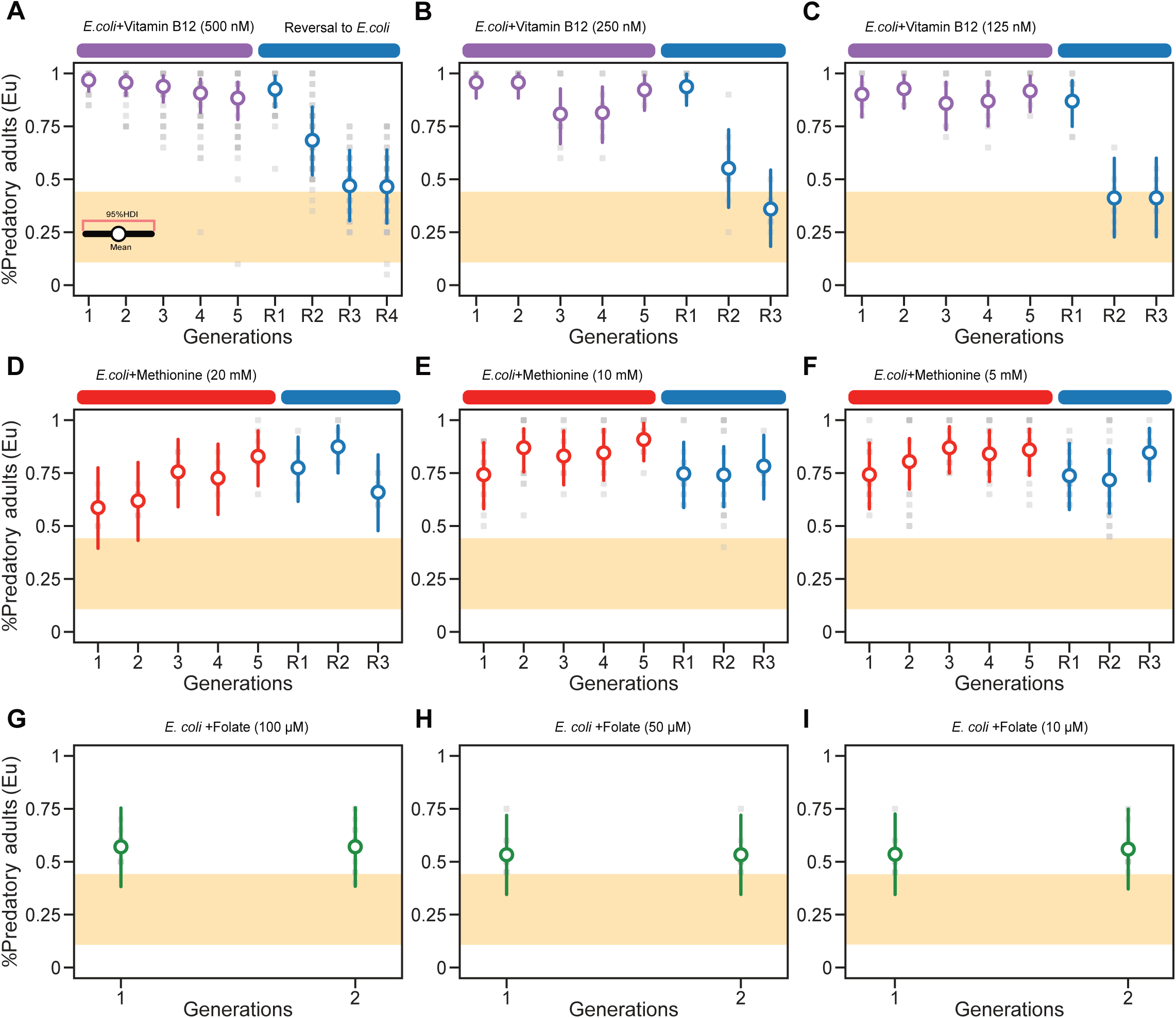
**Memory of the induced predatory mouth form after supplementation with different concentrations of vitamin B12, methionine and folate.** (A-C) Concentration-dependent effect of vitamin B12 on memory transmission. (A) Mean probability of predatory mouth-form after a 5-generation exposure to 500 nM vitamin B12 shows Eu memory for two generations. Further reduction of vitamin B12 concentration to (B) 250 nM, (C) 125 nM, shows a single generation of Eu memory on un-supplemented *E. coli* diets. (D-F) Mean probability of the predatory mouth form on methionine-supplemented *E. coli* plates. (G-I) Mean probability of the predatory mouth form on folate-supplemented *E. coli* plates. See also Figure S4.

However, the response during reversal only lasted for two generations, with all following generations being down to baseline Eu frequencies (0.841 ≤HDI(θ_R1_)≤ 0.987, 0.52 ≤HDI(θ_R2_)≤ 0.842, and 0.305 ≤HDI(θ_R3_)≤ 0.637) (Figures 4A and S4A). Similarly, supplementation experiments using 250 nM and 125 nM vitamin B12, resulted in an immediate Eu mouth-form induction, but reversal experiments after 5 or 10 generations resulted in an even shorter memory that lasted only for one generation (Figures 4B, 4C, S4B and S4C). When we reduced the concentration of vitamin B12 even further and supplemented with 50 nM, 10 nM, 5 nM and 2 nM vitamin B12, all four of these concentrations still resulted in a strong induction of the Eu mouth-form (Figures S4D-S4G). In contrast, worms lost their memory already in the F5R2 generation (for 50nM, 0.276 ≤HDI(θ_R2_)≤ 0.655, for 10 nM, 0.238 ≤HDI(θ_R2_)≤ 0.612, and for 5 nM, 0.27 ≤HDI(θ_R2_)≤ 0.645) (Figures S4D-S4G). Thus, the reduction of vitamin B12 concentration in supplementation experiments results in the loss of TEI of the predatory mouth form, while the induction of the Eu morph remains high. Therefore, TEI of the predatory mouth form requires a higher vitamin B12 concentration than the initial induction of the Eu mouth form.

### Transgenerational inheritance of the predatory morph require methionine but not folate

Next, we used supplementation experiments with two major metabolites in the one-carbon cycle (Figure 3A). First, we used 10 μM, 50 μM and 100 μM of folate (vitamin B9) for supplementation of standard *E. coli* plates and found no credible induction of the predatory mouth form (e.g., for 100 μM, 0.384 ≤HDI(θ_F2_)≤ 0.755, Figures 4G-4I). In contrast, methionine supplementation resulted in the induction of the predatory mouth form and subsequent TEI; however, with different dynamics than what is observed after vitamin B12 supplementation. Specifically, supplementation with 5 mM, 10 mM and 20 mM methionine resulted in the induction of the Eu mouth form after several generations of exposure (Figures 4D-4F). Note that this supplementation does not result in 100% Eu animals similar to supplementation with lower concentrations of vitamin B12 (Figure 4). However, reversal after five generations of exposure to methionine indicates a TEI of the predatory mouth form that lasts for three generations (Figures 4D-4F). Thus, vitamin B12 likely exerts its function through methionine in the one-carbon cycle.

### Vitamin B12 causes elevated nutrient provisioning

It is important to determine what type of changes act downstream of the original stimulus to cause the inherited effect. This is of particular importance given the different vitamin B12 concentrations necessary to induce the predatory mouth form and its TEI. In principle, a multigenerational vitamin B12-rich diet could induce a plethora of transcriptional changes in the worm. Therefore, we extended our gene expression analysis to cultures after reversal and compared transcriptional profiles of naïve, F3, F4, F5 and F10 generations on *Novosphingobium* as well as F3R2, F4R2, F5R2 and F10R2 generations after reversal to *E. coli* (Figure 5A). We used FnR2 rather than FnR1 generations to rule out any effect of the kanamycin treatments in the FnR1 generation. We found a consistently strong increase in vitellogenin expression after *Novosphingobium* exposure with multiple vitellogenin genes remaining at higher expression levels in comparison to naïve animals (FDR-corrected p value <0.01) (Figures 2E and 5B). Specifically, *P. pacificus* shows a more complex vitellogenin composition when compared with *C. elegans*. While the *C. elegans* N2 genome contains six vitellogenin genes, *P. pacificus* RSC011 has 9 genes all of which are most closely related to *Cel-vit-6* (Figure 5C). We therefore named these genes *Ppa-vit-6-A* to *Ppa-vit-6-I* based on their location along the chromosome (Figures 5C and 5D). While more similar to one another in sequence than the *Cel-vit* genes, the divergence between the *Ppa-vit* genes still allows the distinction of individual genes in gene expression analysis (Figure 5B). We found that 8 of the 9 vitellogenin genes are upregulated upon *Novosphingobium* exposure, four of which show an increased expression of at least eightfold in the F10 generation (Figure 5B, Data S3). A similar increase in expression was seen in all other tested generations on *Novosphingobium.* After reversal to *E. coli* the majority of vitellogenin genes were less expressed than on *Novosphingobium*; however, expression remained substantially higher than in naïve animals that had never been exposed to *Novosphingobium* (Figure 5B). Thus, exposure to a multigenerational *Novosphingobium* diet and subsequent reversal to *E. coli* cause elevated vitellogenin transcription resulting in increased nutritional provisioning.

**Figure 5.**
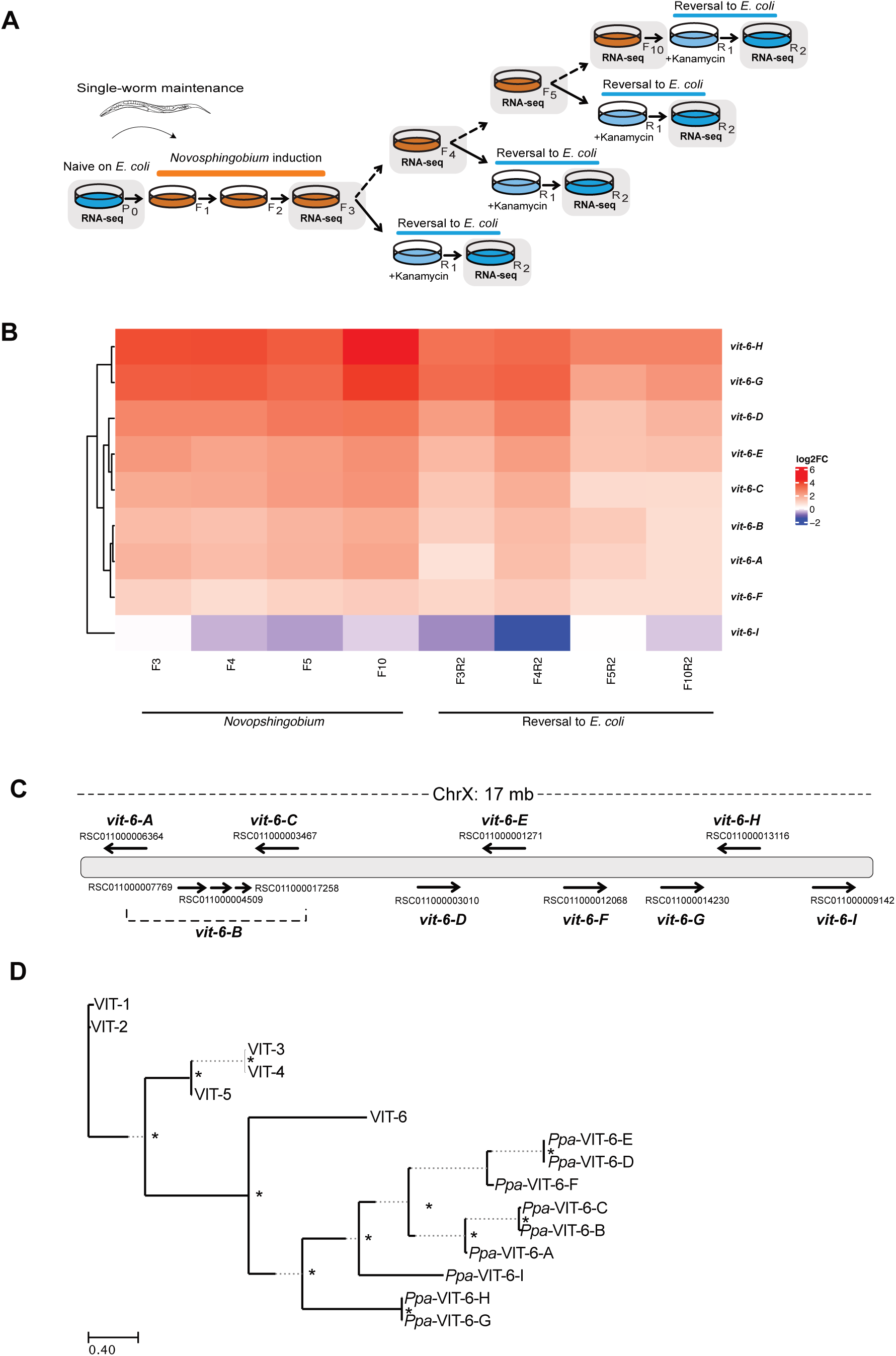
**Vitellogenin gene organization, phylogeny and expression changes in response to dietary switch experiments.** (A) Graphical representation of the experiment. Generations highlighted in grey indicate worm populations sent for RNA sequencing. (B) Vitellogenin expression of all 9 genes under different wildtype conditions: Log_2_FoldChange (log_2_FC) values are shown for worms on a *Novosphingobium* diet (F3, F4, F5, F10) and reversal to *E. coli* (F3R2, F4R2, F5R2, F10R2). Naïve animals on *E. coli* (P0) were used as reference to generate log_2_FC values. (C) *P. pacificus* RSC011 has 9 vitellogenin genes all located on the X chromosome distributed over a 17 Mb range. These genes are named *Ppa-vit-6-A* to *Ppa-vit-6-I* based on their location along the chromosome. Note that there are three transcripts that encodes for *Ppa-vit-6-B*. (D) Phylogeny of the *P. pacificus* and *C. elegans* vitellogenin genes. In *C. elegans*, six vitellogenin genes show higher sequence divergence with the *Cel-vit-6* gene being the most diverse gene copy. Note that it is *Cel-vit-6* that is most similar to the individual genes in *P. pacificus* and also other nematodes. Nodes with bootstrap values of ≥90 are labelled with asterisks (*).

### Receptor-mediated endocytosis (*rme-2*) mutants have no memory

The correlation between elevated vitellogenin expression and the formation of the predatory mouth-form during and after *Novosphingobium* exposure suggests a role of nutritional provisioning in transgenerational memory. To provide mechanistic evidence for this hypothesis, we investigated the receptor involved in vitellogenin uptake into the germline. In nematodes, vitellogenin is produced in the intestine and the low-density lipoprotein receptor (LDLR) encoded by *rme-2* (*r*eceptor-*m*ediated *e*ndocytosis protein) was shown to represent the single protein involved in vitellogenin uptake into the germline.^19,20^ In *C. elegans*, *rme-2* mutant embryos contain no detectable yolk and the brood size of homozygous mutant animals is strongly reduced.^19^ In *P. pacificus*, there is a 1:1 ortholog *Ppa-*RME-2 with a 55% amino acid sequence similarity (Figure 6A). Strikingly, we had isolated three mutant alleles of *Ppa-rme-2* in our EMS mutagenesis screen for predatory mouth-form Transgenerational-inheritance-defective (Tid) mutants reported previously (Figure 6A; Table S2).^14^ This finding would be consistent with a role of vitellogenin in memory formation.

**Figure 6.**
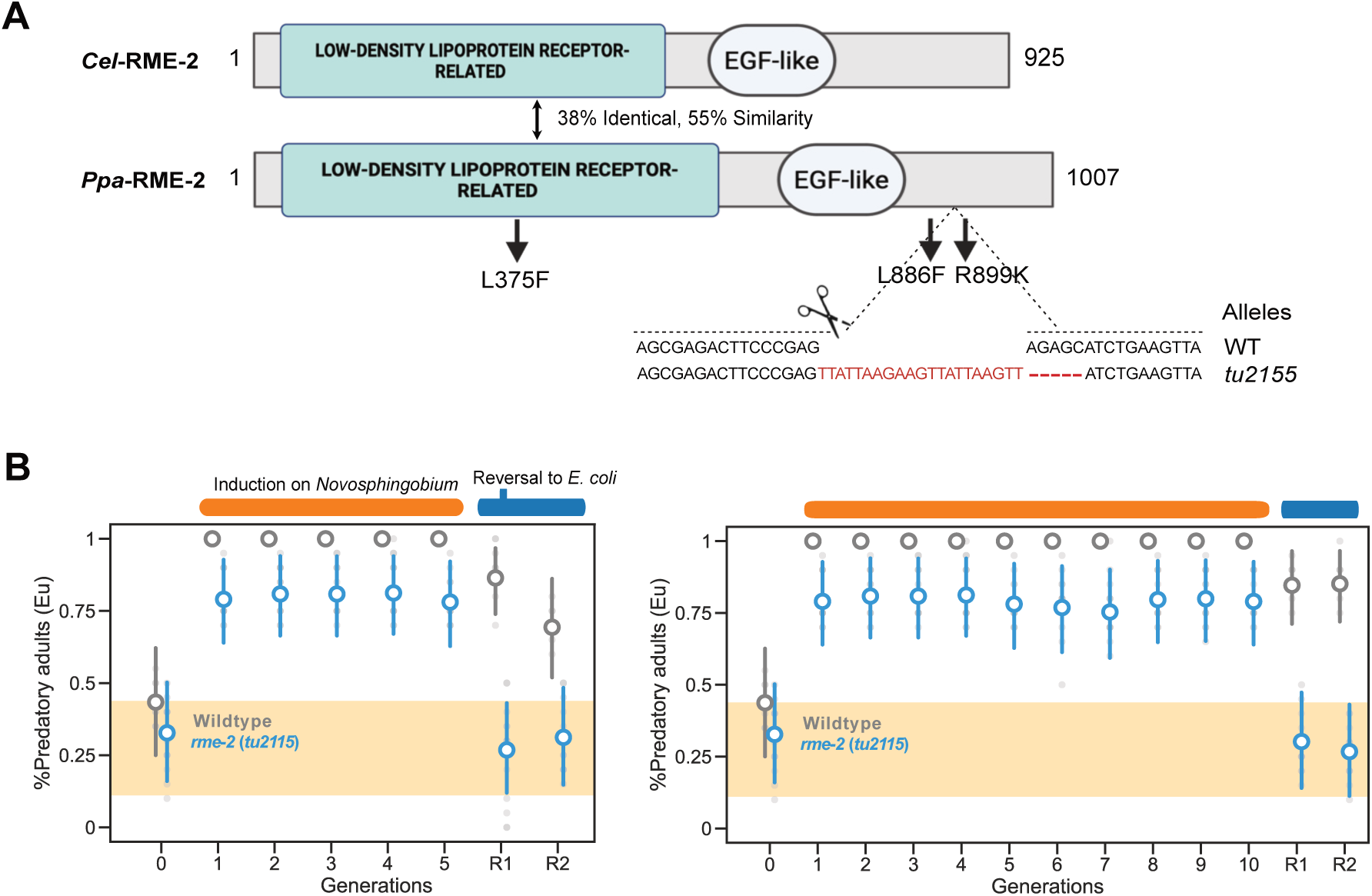
**The low-density lipoprotein receptor related protein RME-2 is required for vitamin B12 transmission and transgenerational inheritance of the predatory morph** (A) Domain architecture of *C. elegans* and *P. pacificus* RME-2, sequence similarity and position of molecular lesions in three *Ppa-rme-2* mutants are highlighted. Three alleles isolated in a forward genetic EMS mutagenesis screen carry point mutations resulting in amino acid changes as indicated. We designed a sgRNA in the C-terminal region where two of the original EMS alleles carry their mutation. The frameshift allele *tu2215* was used for further analysis. (B) Mean probability of the predatory mouth form of *Ppa-rme-2(tu2215)* after a *Novosphingobium* exposure of 5 (left) and 10 (right) generations.

To rule out the possibility that other compensatory mutations after EMS mutagenesis are involved in the observed Tid phenotype in the mutants mentioned above, we generated an additional knock-out mutation by CRISPR engineering. The *Ppa-rme-2(tu2155)* allele has a 16 bp insertion that results in a premature stop codon (Figure 6A). When we tested the *Ppa-rme-2(tu2155)* allele in a 5 generation *Novosphingobium* assay, we observed an incomplete induction of the Eu morph for all five generations of *Novosphingobium* exposures (0.64 ≤HDI(θ_F1-*tu2115*_)≤ 0.928, 0.664 ≤HDI(θ_F2_)≤ 0.94, and 0.628 ≤HDI(θ_F5_)≤ 0.921) (Figure 6B). This result indicates that in the absence of RME-2 and vitellogenin uptake into the germline, the predatory mouth-form can still be induced, which is likely due to other *Novosphingobium*-derived factors as previously indicated (Figure 2C). In contrast, after the reversal to *E. coli*, *Ppa-rme-2(tu2155)* did not show transgenerational memory of the predatory morph (0.12 ≤HDI(θ_F5R1-*tu2115*_)≤ 0.431, 0.147 ≤HDI(θ_F5R2_)≤ 0.484) (Figure 6B). Thus, vitellogenin is involved in the formation of transgenerational memory of the predatory mouth-form in *P. pacificus* indicating a role for nutrient provisioning in this process.

## DISCUSSION

This work demonstrates that a food-derived vitamin can cause transgenerational memory of a morphological trait and its associated behavior over multiple generations. The identification of vitamin B12 as inducing stimulus for the TEI of the predatory mouth form in *P. pacificus* after multigenerational exposure to *Novosphingobium* indicates the significance of bacterial metabolites for animal development, growth and behavior and provides links to vitamin B12 deficiency in humans. Vitamin B12 (cobalamin) is the largest and most complex vitamin.^18,21^ Exclusively synthesized by bacteria, vitamin B12 is essential for animals including humans. However, not all bacteria produce vitamin B12 and the standard *C. elegans*/*P. pacificus* food source, *E. coli* OP50, is a non-vitamin B12 producer.

Therefore, under standard laboratory growth conditions, vitamin B12 is one of the growth-limiting factors for *E. coli* bacteria and worms, and both obtain vitamin B12 exclusively from the tryptone in the agar plates.^16^ Consistently, previous work indicated that vitamin B12 supplementation or a vitamin B12-rich *Comamonas* or *Novosphingobium* diet affect several *C. elegans* life history traits and killing efficiency in *P. pacificus* .^15,17,22,23^

Bacterial and animal requirement for vitamin B12 is low. For example, for *E. coli* it was estimated that only 20 cobalamin molecules per cell are sufficient to support growth.^24^ In humans, the total body content of vitamin B12 is estimated to be 1-5 mg with a daily requirement of 2-3 *μ*g.^25,26^ In *C. elegans,* studies by Bito and co-worker have shown that culture of worms under strict vitamin B12-deficient conditions will result in a gradual decrease of the vitamin B12 content over five generations.^16^ In the fifth generation, *C. elegans* had only 4% of vitamin B12 relative to worms growing under standard conditions and started to show a loss of fertility and reduced lifespan at these reduced concentrations.^16^ These findings are consistent with three of our observations; i) the multigenerational exposure to a *Novosphingobium* diet being necessary to induce TEI of the predatory mouth form, ii) the capability of vitamin B12 supplementation to induce TEI of this phenotype, and most importantly iii) the concentration-dependent effect of vitamin B12 supplementation. It is important to note that most studies in nematodes used un-physiological amounts of vitamin B12 supplementation, usually 1,000 or 500 nM. But vitamin B12 is not directly taken up by the worms from the agar and instead is ingested through the *E. coli* diet. Given that the direct measurement of vitamin B12 still represents a major challenge, the exact amount of vitamin B12 in *C. elegans* or *P. pacificus* after a *Comamonas* or *Novosphingobium* diet is currently unknown, a limitation that is common to all studies on vitamin B12.^27^

In *P. pacificus*, the inherited effect of vitamin B12 acts through vitellogenin. Vitellogenins are a family of yolk proteins representing the most abundant proteins in oviparous animals. Previous work in *C. elegans* had indicated a role of vitellogenins for post-embryonic development and fertility.^20^ Increased vitellogenin provisioning is associated with several post-embryonic phenotypic alterations, i.e. advanced maternal age. Specifically, young *C. elegans* mothers provide less embryonic vitellogenin resulting in early-hatched larvae being smaller and slower to reach adulthood. In contrast, older mothers provide more vitellogenin resulting in larger larvae that are also more starvation resistant, indicating an intergenerational signal that mediates the influence of parental physiology on progeny.^20,28^ Our work extends these findings indicating that nutrient provisioning affects organismal physiology and behavior involving transgenerational inheritance and requires epigenetic factors. This multigenerational effect highlights the significance of exceptions to the Weissmann barrier and suggests vitellogenin transmission from somatic tissues into the germline to have important consequences for development and evolution.

## Acknowledgments

We would like to thank the entire support team and technicians of the Sommer lab in particular Heike Hausmann for assisting in freezing nematode cultures. We are also grateful to Drs. Adrian Streit and Catia Igreja for discussions and helpful comments on the manuscript. The work was funded by the Max Planck Society.

## Author contributions

Conceptualization, S.P.Q. and R.J.S.; Methodology, S.P.Q., A.K., C.R. and R.J.S; Formal Analysis, S.P.Q., A.K.; Investigation, S.P.Q.; Resources, R.Z. and H.W.; Writing – Original Draft, S.P.Q. and R.J.S.; Writing – Review & Editing, S.P.Q., A.K., C.R. and R.J.S.; Visualization, S.P.Q. and A.K.; Funding Acquisition, R.J.S; Supervision, R.J.S.

## Supplemental Information

**Supplemental Figures. Figures S1–S4 and Table S1-S2.**

Data S1. Differential gene expression analysis and enriched gene sets after exposure to *Novosphingobium* for three and five generations. Related to Figure 2 and STAR methods.

(A and B) Upregulated and downregulated gene lists after three and five generations on *Novosphingobium.* (C and D) Lists of Pfam-domains enriched in the differentially expressed gene sets after three and five generations on *Novosphingobium*.

Data S2. Differential gene expression analysis and enriched gene sets after exposure to *E. coli* supplemented with vitamin B12 for three and five generations. Related to Figure 2 and STAR methods

(A and B) Upregulated and downregulated gene lists after three and five generations of vitamin B12 supplementation. (C and D) Lists of Pfam-domains enriched in the differentially expressed gene sets after three and five generations on *Novosphingobium*.

Data S3. List of differentially expressed genes upon *Novosphingobium* exposure and reversal to *E. coli.* Related to Figure 5.

## STAR Methods

### Resource availability Lead contact

Further information and requests for resources and reagents should be directed to and will be fulfilled by the lead contact Ralf J. Sommer.

### Materials availability

*P. pacificus* strains and bacterial isolates generated in this work are freely available through the Lead Contact.

### Data and code availability

All the code and data for Bayesian analysis are available at https://github.com/shielapearl18/Bayesian-estimates-of-mouth-form-responses-in-vitamin-B12-enriched-environment.

Sequencing data that was generated for the current study has been submitted to the European Nucleotide Archive under the project accession PRJEB74486.

### Experimental model and subject details

The ancestral *P. pacificus* RSC011 isolate used in this study was frozen within the first 5-10 generations after original isolation from Coteau Kerveguen on La Réunion to minimize domestication and thereby, facilitate the investigation of diet-induced plasticity and transgenerational epigenetic inheritance. Nematodes were grown under standard nematode growth conditions on NGM plates seeded with either *Escherichia coli* OP50 or *Novosphingobium* L76 and maintained at 20°C. The RSC011 stock was not subjected to bleaching, starvation, extreme temperature fluctuations as they are known to influence mouth-form ratios.

### Bacterial strains conditions

All bacterial strains and mutants were grown overnight in LB (Lysogeny broth) supplemented with 50 μg/ml kanamycin where required to initiate food reversal. Bacteria were grown at 30 °C or 37 °C depending on the species and 6 cm nematode growth medium (NGM) plates were seeded with 300 μl bacterial overnight cultures and were incubated for 2 days.

### Method details

#### Nematode culture and dietary reversal experiments

Overnight cultures of *E. coli* OP50 and *Novosphingobium* L76 were spread to NGM plates and incubated at room temperature for 2 days. The *P. pacificus* RSC011 strain is preferentially St (20-40% Eu) on a standard *E. coli* OP50 diet.

*Novosphingobium* exposure is performed by picking single J4 larvae onto a *Novosphingobium* L76 diet. J4 worms were initially left for at least an hour on an initial *Novosphingobium* diet to reduce traces of *E. coli* OP50 before transferring to final F1 *Novosphingobium* plates. Reversal experiments to an *E. coli* OP50 diet were performed after exposing of RSC011 worms for various numbers of generations, i.e. 1, 3, 5, 10 or 15 generations on *Novosphingobium* L76. Worms were initially transferred from *Novosphingobium* L76 to NGM plates supplemented with 50 μg/ml of kanamycin for one generation, then back to *E. coli* OP50-seeded NGM plates for subsequent generations. Exposure to kanamycin for a single generation to eliminate traces of *Novosphingobium* did not affect mouth-form response.^14^

#### Mouth-form phenotyping

Mouth-from phenotyping was performed using Zeiss Discovery V.20 stereo microscope (X150 magnification) by observing the nematode buccal cavity based on mouth-form identities previously described.^8^ Final mouth-form frequencies are the mean of at least 20 replicates, each assaying 20 animals from a single plate.

#### Statistical analysis

To estimate the probability of developing the probability of developing the Eu mouth form in *P. pacificus* (**θ**), we constructed a hierarchical Bayesian model as previously described^14^

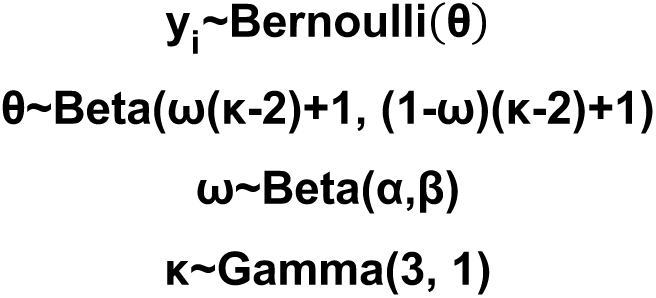

where ***θ*** is calculated for each replicate and hyperparameters μ and κ link the biological replicates for a given generation under a given experimental condition.^29^ The model was fitted to the laboratory measurements using PyMC^30^ in Python 3.11 with Numpy 1.25.2.^31^ The mean highest density interval (HDI) for **θ** for a group of observations was used to visualize the inferred probability of developing the Eu mouth form. We ensured that the estimated **θ** values were stable using common convergence diagnostics with effective sample size (ESS) **≥**10000.^32^

#### CRISPR/Cas9-Induced Mutations

Mutations in candidate genes were generated using CRISPR/Cas9 following the previously described protocols.^33^ Gene-specific crRNAs and universal trans-activating CRISPR RNA (tracrRNA) were obtained from Integrated DNA Technologies. Equal volumes of (5 μl) of each 100 μM stock were mixed and denatured at 95 °C for 5 min, then allowed to anneal at room temperature. Cas9 endonuclease (New England Biolab) was added to the hybridized product and incubated at room temperature for 5 min. TE buffer was used to adjust to final concentrations of 18.1 μM for the sgRNA and 2.5 μM for Cas9. The resulting mixture was microinjected into the germline of 40-50 *P. pacificus* RSC011 young adults.

Eggs from injected P0s were collected up to 16 hours post-injection. After hatching and two days of growth, the F1 progeny was segregated onto individual plates until they had fully-developed and laid eggs sufficiently. F1 genotypes were screened using Sanger sequencing and mutations were identified prior to isolation of homozygous mutants. Details of sgRNAs and primers used in this study are provided in the supplementary table. Evidence for null alleles was based on frameshift mutations causing premature stop codons in protein coding sequences.

#### Metabolite supplementation assays

Methylcobalamin (Vitamin B12 CAS No. 13422–55–4), L-methionine (CAS No. 63– 68–3) and folate (CAS No. 59-30-3) were purchased from Sigma and dissolved in water at the highest possible soluble concentrations to prepare stock solution. All stocks were prepared fresh before use in each experiment. Metabolite solutions were mixed with NGM agar at the required concentration just before pouring to 6 cm plates. Plates were allowed to dry at room temperature for two days and then spotted with *E. coli* OP50. We first tested different concentrations of vitamin B12 and found the strongest and most reliable transgenerational epigenetic memory effect with a concentration of 1000 nM and 1500 nM, which is most likely un-physiological. Similarly in *C. elegans*, dose-dependent effects have been seen for vitamin B_12_.^17^ Different concentrations for methionine and folate supplementation were based on previous studies.^15,34^

#### RNA sequencing and data analysis

Mixed-staged RSC011 worms, reared on *Novosphingobium* and *E. coli* OP50 supplemented with vitamin B12, were harvested from three NGM plates. Worms were pelleted for RNA extraction after 3 and 5 generations of dietary exposure. Total RNA was isolated using Direct-Zol RNA Mini prep kit (Zymo Research) following the manufacturer’s instructions. The RNA-seq library preparation and sequencing were performed by the company Novogene. Raw reads were aligned to the *P. pacificus* RSC011 reference genome using Hisat2^35^ (version 2.1.0), and read counts were quantified with featureCounts^36^ based on the RSC011 gene annotations. For the analysis of vitellogenenin expression, we combined raw counts of the three transcripts (RSC011000007769, RSC011000004509 and RSC011000017258) into a single *vit-6-B* count. Differential gene expression analysis was performed in R (version 4.0.3) using DESeq2 (version 1.18.1).^37^ Genes with an FDR-corrected *p* value < 0.01 and fold change cutoff of two were considered as significantly differentially expressed. Tests for overrepresentation of protein domains in sets of differentially expressed genes were performed using a Fisher’s exact test in R. We used the FDR method as implemented in the p.adjust function in R to correct for multiple testing and only retained results with adjusted P-value < 0.01. Overlap between gene sets affected by vitamin B12 supplementation and *Novosphingobium* exposure was evaluated using Fisher’s exact test implemented in the GeneOverlap package (version 1.24.0).^38^

#### Phylogenetic analysis

One- to- one orthologous genes between *P. pacificus* and *C. elegans* was identified from BLASTP searches and were validated by phylogenetic analysis. Proteins were aligned using Clustal Omega^39^ and the output FASTA file was uploaded to the IQ-TREE tool.^40,41^ Analysis was performed under default settings using the auto substitution model, and 1000 bootstrap alignments were calculated with ultrafast setting. FigTree was used to visualized the resulting phylogenetic tree (http://tree.bio.ed.ac.uk/software/figtree/).

## References

1. Painter, R.C., Osmond, C., Gluckman, P., Hanson, M., Phillips, D.I.W., and Roseboom, T.J. (2008). Transgenerational effects of prenatal exposure to the Dutch famine on neonatal adiposity and health in later life. BJOG 115, 1243–1249.

2. Song, S., Wang, W., and Hu, P. (2009). Famine, death, and madness: Schizophrenia in early adulthood after prenatal exposure to the Chinese Great Leap Forward Famine. Soc. Sci. Med. 68, 1315–1321.

3. Baugh, L.R., and Day, T. (2020). Nongenetic inheritance and multigenerational plasticity in the nematode *C . elegans*. Elife 9, 1–13.

4. Frolows, N., and Ashe, A. (2021). Small RNAs and chromatin in the multigenerational epigenetic landscape of *Caenorhabditis elegans*. Philos. Trans. R. Soc. B Biol. Sci. 376, 1–12.

5. Chen, X., and Rechavi, O. (2022). Plant and animal small RNA communications between cells and organisms. Nat. Rev. Mol. Cell Biol. 23, 185–203.

6. Santilli, F., and Boskovic, A. (2023). Mechanisms of transgenerational epigenetic inheritance: lessons from animal model organisms. Curr. Opin. Genet. Dev. 79, 1–8.

7. Bento, G., Ogawa, A., and Sommer, R.J. (2010). Co-option of the hormone-signalling module dafachronic acid-DAF-12 in nematode evolution. Nature 466, 494–497.

8. Ragsdale, E.J., Müller, M.R., Rödelsperger, C., and Sommer, R.J. (2013). A developmental switch coupled to the evolution of plasticity acts through a sulfatase. Cell 155, 922–933.

9. Lightfoot, J.W., Wilecki, M., Rödelsperger, C., Moreno, E., Susoy, V., Witte, H., and Sommer, R.J. (2019). Small peptide-mediated self-recognition prevents cannibalism in predatory nematodes. Science. 364, 86–89.

10. Ptashne, M. (1986). A genetic switch: Gene control and phage. J. Med. Genet. 24, 789–790.

11. Balázsi, G., Van Oudenaarden, A., and Collins, J.J. (2011). Cellular decision making and biological noise: From microbes to mammals. Cell 144, 910–925.

12. Sommer, R.J. (2020). Phenotypic plasticity: From theory and genetics to current and future challenges. Genetics 215, 1–13.

13. Dardiry, M., Piskobulu, V., Kalirad, A., and Sommer, R.J. (2023). Experimental and theoretical support for costs of plasticity and phenotype in a nematode cannibalistic trait. Evol. Lett. 7, 48–57.

14. Quiobe, S.P., Kalirad, A., Roeseler, W., Witte, H., Wang, Y., Roedelsperger, C., and Sommer, R.J. (2025). EBAX-1/ZSWIM8 destabilizes miRNAs resulting in transgenerational memory of a predatory trait. Sci. Adv. 11, 1–17.

15. Akduman, N., Lightfoot, J.W., Röseler, W., Witte, H., Lo, W.S., Rödelsperger, C., and Sommer, R.J. (2020). Bacterial vitamin B12 production enhances nematode predatory behavior. ISME J. 14, 1494–1507.

16. Bito, T., Matsunaga, Y., Yabuta, Y., Kawano, T., and Watanabe, F. (2013). Vitamin B12 deficiency in *Caenorhabditis elegans* results in loss of fertility, extended life cycle, and reduced lifespan. FEBS Open Bio 3, 112–117.

17. Watson, E., Macneil, L.T., Ritter, A.D., Yilmaz, L.S., Rosebrock, A.P., Caudy, A.A., and Walhout, A.J.M. (2014). Interspecies systems biology uncovers metabolites affecting *C. elegans* gene expression and life history traits. Cell 156, 759–770.

18. Litwack, G. (2022). Vitamins and Hormones (Academic Press: Cambridge, MA).

19. Grant, B., and Hirsh, D. (1999). Receptor-mediated endocytosis in the *Caenorhabditis elegans* oocyte. Mol. Biol. Cell 10, 4311–4326.

20. Perez, M.F., and Lehner, B. (2019). Vitellogenins - Yolk Gene Function and Regulation in *Caenorhabditis elegans*. Front. Physiol. 10, 1–17.

21. Eschenmoser, A. (2011). Etiology of potentially primordial biomolecular structures: From vitamin B12 to the nucleic acids and an inquiry into the chemistry of life’s origin: A retrospective. Angew. Chemie - Int. Ed. 50, 12412–12472.

22. Watson, E., MacNeil, L.T., Arda, H.E., Zhu, L.J., and Walhout, A.J.M. (2013). Integration of metabolic and gene regulatory networks modulates the *C. elegans* dietary response. Cell 153, 253–266.

23. MacNeil, L.T., Watson, E., Arda, H.E., Zhu, L.J., and Walhout, A.J.M. (2013). Diet-induced developmental acceleration independent of TOR and insulin in *C. elegans*. Cell 153, 240–252.

24. Di Girolamo, P.M., Kadner, R.J., and Bradbeer, C. (1971). Isolation of Vitamin B12 Transport Mutants of *Escherichia coli*. J. Bacteriol. 106, 751–757.

25. Doets, E.L., In’t Veld, P.H., Szczecińska, A., Dhonukshe-Rutten, R.A.M., Cavelaars, A.E.J.M., Van ’t Veer, P., Brzozowska, A., and De Groot, L.C.P.G.M. (2013). Systematic review on daily Vitamin B12 losses and bioavailability for deriving recommendations on Vitamin B12 intake with the factorial approach. Ann. Nutr. Metab. 62, 311–322.

26. Institute of Medicine (1998). Vitamin B12. In Dietary Reference Intakes for Thiamin, Riboflavin, Niacin, Vitamin B6, Folate, Vitamin B12, Pantothenic Acid, Biotin and Choline. (Washington DC:National Academies Press), pp. 306–348.

27. Lawrence, A.D., Nemoto-Smith, E., Deery, E., Baker, J.A., Schroeder, S., Brown, D.G., Tullet, J.M.A., Howard, M.J., Brown, I.R., Smith, A.G., et al. (2018). Construction of Fluorescent Analogs to Follow the Uptake and Distribution of Cobalamin (Vitamin B12) in Bacteria, Worms, and Plants. Cell Chem. Biol. 25, 941–951.e6.

28. Perez, M.F., Francesconi, M., Hidalgo-Carcedo, C., and Lehner, B. (2017). Maternal age generates phenotypic variation in *Caenorhabditis elegans*. Nature 552, 106–109.

29. Kruschke, J.K. (2014). Doing Bayesian data analysis: A tutorial with R, JAGS, and Stan, second edition (Elsevier Science).

30. Abril-Pla, O., Andreani, V., Carroll, C., Dong, L., Fonnesbeck, C.J., Kochurov, M., Kumar, R., Lao, J., Luhmann, C.C., Martin, O.A., et al. (2023). PyMC: a modern, and comprehensive probabilistic programming framework in Python. PeerJ Comput. Sci. 9, 1–35.

31. Harris, C.R., Millman, K.J., van der Walt, S.J., Gommers, R., Virtanen, P., Cournapeau, D., Wieser, E., Taylor, J., Berg, S., Smith, N.J., et al. (2020). Array programming with NumPy. Nature 585, 357–362.

32. Kruschke, J.K. (2021). Bayesian Analysis Reporting Guidelines. Nat. Hum. Behav. 5, 1282–1291.

33. Witte, H., Moreno, E., Rödelsperger, C., Kim, J., Kim, J.S., Streit, A., and Sommer, R.J. (2014). Gene inactivation using the CRISPR/Cas9 systemin the nematode *Pristionchus pacificus*. Dev. Genes Evol. 225, 55–62.

34. Virk, B., Correia, G., Dixon, D.P., Feyst, I., Jia, J., Oberleitner, N., Briggs, Z., Hodge, E., Edwards, R., Ward, J., et al. (2012). Excessive folate synthesis limits lifespan in the *C. elegans*: *E. coli* aging model. BMC Biol. 10.

35. Kim, D., Langmead, B., and Salzberg, S.L. (2015). HISAT: A fast spliced aligner with low memory requirements. Nat. Methods 12, 357–360.

36. Liao, Y., Smyth, G.K., and Shi, W. (2014). FeatureCounts: An efficient general purpose program for assigning sequence reads to genomic features. Bioinformatics 30, 923–930.

37. Love, M.I., Huber, W., and Anders, S. (2014). Moderated estimation of fold change and dispersion for RNA-seq data with DESeq2. Genome Biol. 15, 1–21.

38. Shen, L. (2023). GeneOverlap : An R package to test and visualize gene overlaps. https://bioconductor.org/packages/release/bioc/html/GeneOverlap.html

39. Gabler, F., Nam, S., Till, S., Mirdita, M., Steinegger, M., Söding, J., Lupas, A.N., and Alva, V. (2020). Protein Sequence Analysis Using the MPI Bioinformatics Toolkit. Curr. Protoc. Bioinforma. 72, 1–30.

40. Nguyen, L.T., Schmidt, H.A., Von Haeseler, A., and Minh, B.Q. (2015). IQ-TREE: A fast and effective stochastic algorithm for estimating maximum-likelihood phylogenies. Mol. Biol. Evol. 32, 268–274.

41. Trifinopoulos, J., Nguyen, L.T., von Haeseler, A., and Minh, B.Q. (2016). W-IQ-TREE: a fast online phylogenetic tool for maximum likelihood analysis. Nucleic Acids Res. 44, W232–W235.

