## Supplemental Figures for "Diet-derived vitamin B12 induces transgenerational inheritance of nematode predation through elevated vitellogenin provisioning"

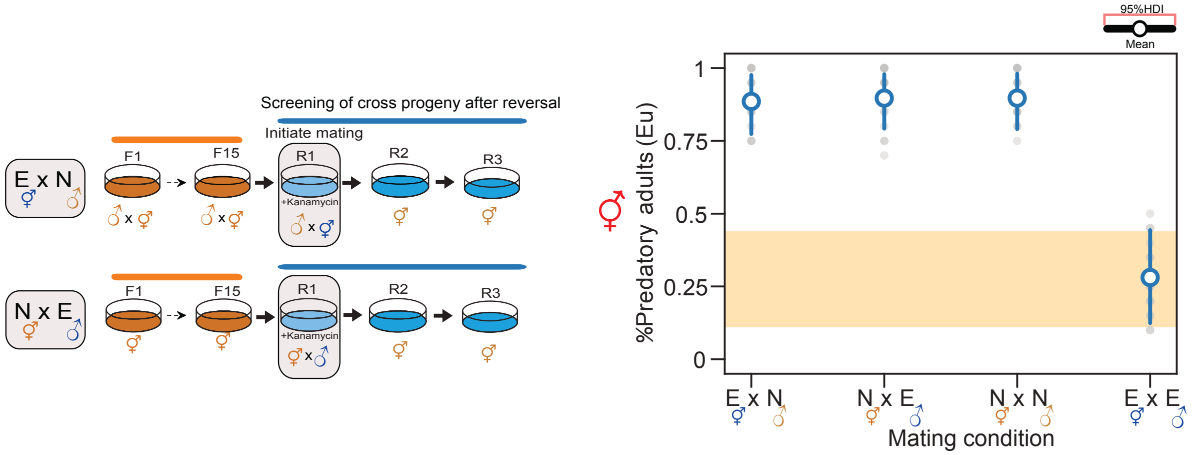


**Figure S1. Mating experimental control conditions. Related to Figure 1.**

Left panel: A schematic representation of the mating experiment. Right panel: Mean probability of the predatory mouth-form during reversal of cross progenies in all different conditions of parental exposures. *Novosphingobium*-exposed males crossed with *Novosphingobium*-exposed hermaphrodites (NxN), and *E. coli*-grown males crossed with *E. coli*-grown hermaphrodites (ExE) were used as controls. Mouth-form screening was performed during the initiation of the mating experiment on the F15R1 generation. N, *Novosphingobium;* E, *E. coli*.

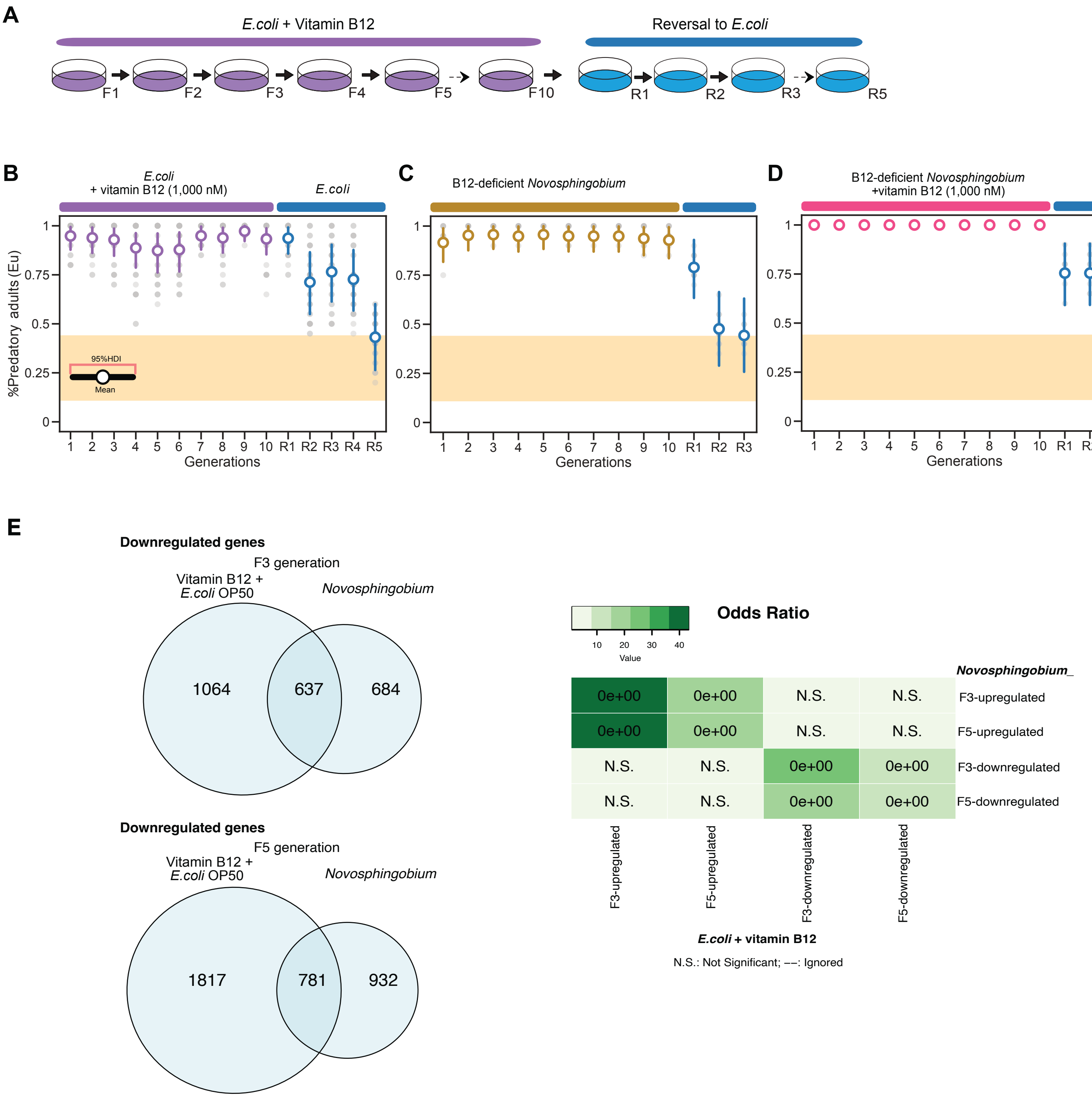


**Figure S2. Induction of the predatory mouth form by vitamin B12- supplemented *E. coli*, *Novosphingobium* vitamin B12-deficient mutant and vitamin B12 rescue. Related to Figure 2.**

(A) Schematic diagram of worms grown on *E. coli* supplemented with vitamin B12 for 10 generations and reversal to un-supplemented *E. coli*. (B) Mean probability of the predatory mouth form after 10 generations of vitamin B12 supplementation with 1,000 nM vitamin B12 and reversal to un-supplemented *E. coli*. (C) Mean probability of the predatory mouth form after 10 generations on a *Novosphingobium* vitamin B12-deficient mutant and reversal to *E. coli*. (D) Mean probability of the predatory mouth form after 10 generations on a *Novosphingobium* vitamin B12-deficient mutant supplemented with exogenous vitamin B12 and reversal to *E. coli*. (E) Overlap of downregulated transcripts during induction on *Novosphingobium* and vitamin B12-supplemented *E. coli* for three and five generations.

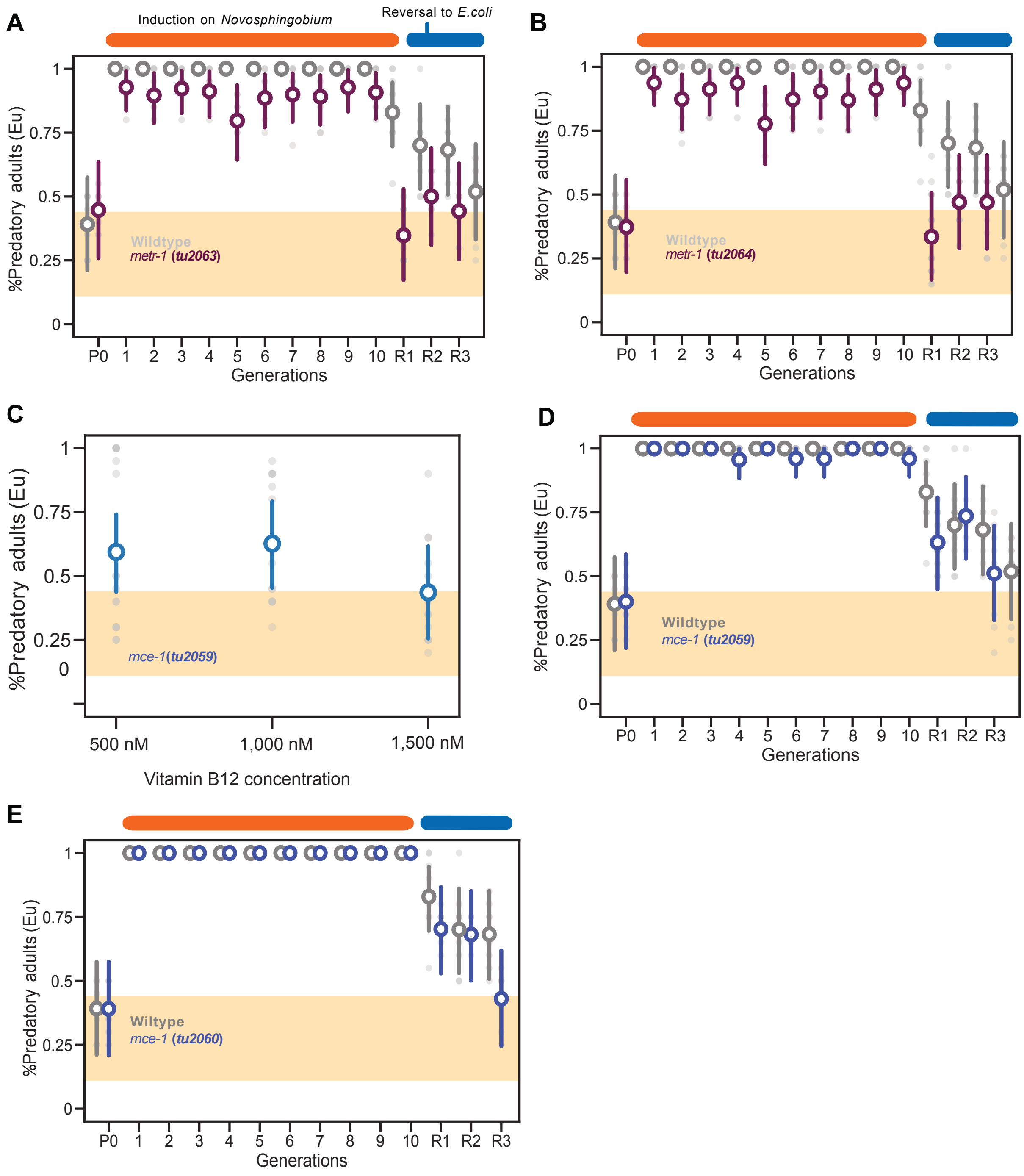


**Figure S3. *Ppa-metr-1* and *Ppa-mce-1* response on *Novosphingobium* and reversal to *E. coli*. Related to Figure 3.**

(A-B) Mean probability of the predatory mouth form in *Ppa-metr-1* mutant animals on *Novosphingobium* after 10 generations of exposure and reversal to *E. coli*. (C) Mean probability of the predatory mouth form in *Ppa-mce-1* mutant animals on vitamin B12-supplemented *E. coli* plates. (D-E) Mean probability of the predatory mouth form in *Ppa-mce-1* mutant animals on *Novosphingobium* after 10 generations of exposure and reversal to *E. coli*. More details about the molecular lesions are listed in Table S1.

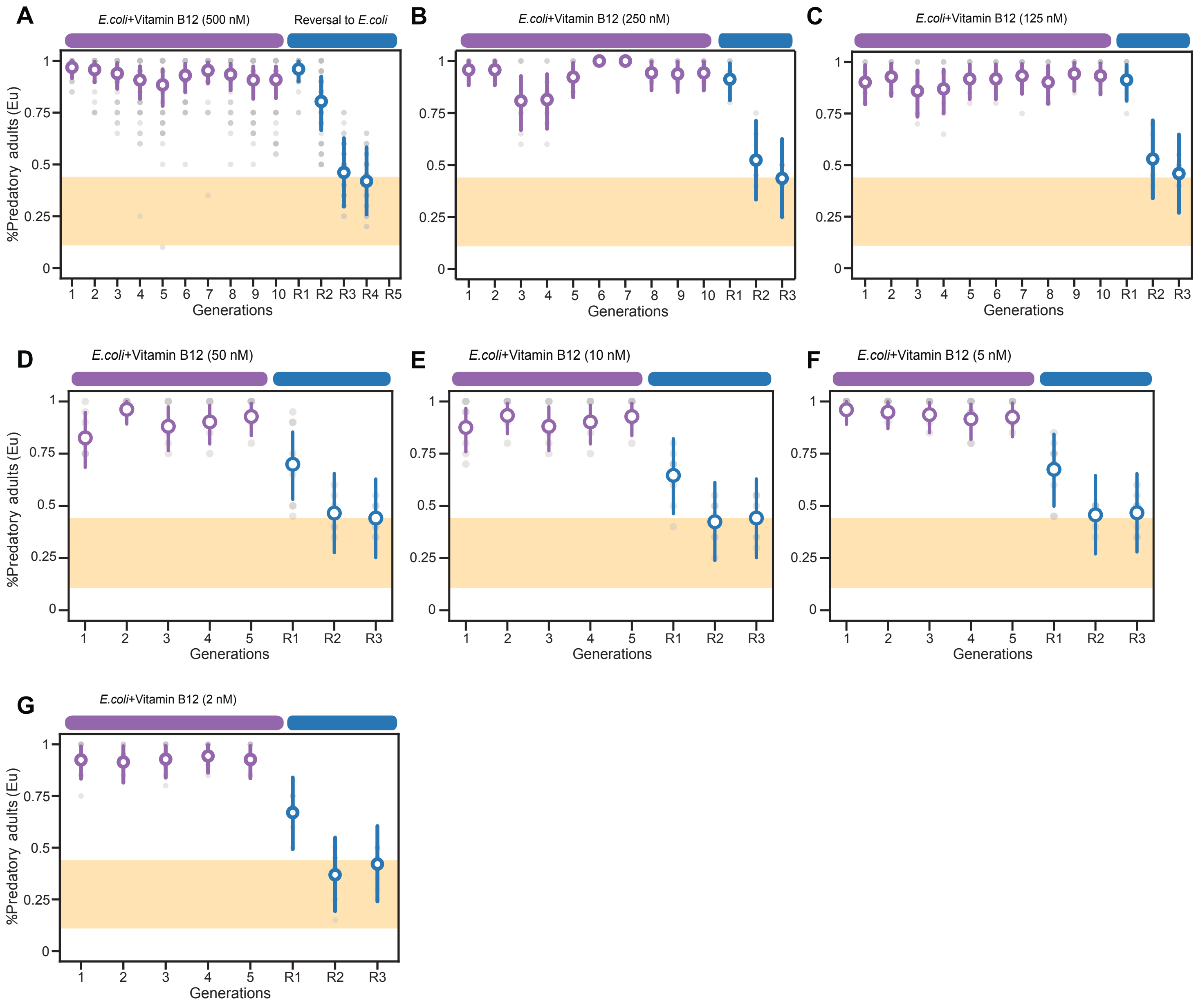


**Figure S4. Vitamin B12 supplementation on the induced predatory mouth form and its maternal effect after reversal on *E. coli*. Related to Figure 4.**

(A-C) Mean probability of predatory mouth-form on 500, 250, 125 nM of vitamin B12 supplementation for 10 generations and subsequent exposure to un-supplemented *E. coli*. (D-F) Mean probability of predatory mouth-form on 50, 10, 5 nM of vitamin B12 supplementation for 5 generations and subsequent exposure to un-supplemented *E. coli*. (G) Mean probability of predatory mouth-form on 2nM of vitamin B12 for 5 generations and exposure to un-supplemented *E. coli*.

| Alleles | Gene ID; *C. elegans* best hit | Molecular lesions | sgRNA(PAM) | Forward primer | Reverse primer |
|---|---|---|---|---|---|
| *tu2063* | RSC011000010081; *metr-1* | 10 bp net insertion | AGAAATGAGACCATTCGTAG(AGG) | GAGAACCACACTCCGATACG | CAACTTGACTACGAGCTATTG |
| *tu2064* | | 55 bp net insertion | | | |
| *tu2059* | RSC011000034399; *mce-1* | 13 bp insertion | CGCCACTCCGGACATCGAGA(AGG) | CGCCAATCTTACCAGAGTGA | CATTGATGTCCTTGACCTAC |
| *tu2060* | | 28 bp deletion | | | |

**Table S1. Molecular lesions of *Ppa-metr-1* and *Ppa-mce-1*. Related to Figure 3 and STAR Methods.**

| Alelle | Gene ID | *C. elegans* 1:1 ortholog | Position | Reference | Variant | Mutation type | Nucleotide substitution |
|---|---|---|---|---|---|---|---|
| | RSC011000018058 | *rme-2* | 12582507 | G | A | nonsynonym [R>K] | AGG>AAG |
| | RSC011000018058 | *rme-2* | 12582467 | C | T | nonsynonym [L>F] | CTT>TTT |
| *tu797* | RSC011000018058 | *rme-2* | 12579999 | C | T | nonsynonym [L>F] | CTC>TTC |

**Table S2. *Ppa-rme-2* alleles isolated from the genetic screen by EMS mutagenesis.**
